# How the tortoise beats the hare: Slow and steady adaptation in structured populations suggests a rugged fitness landscape in bacteria

**DOI:** 10.1101/005793

**Authors:** Joshua R. Nahum, Peter Godfrey-Smith, Brittany N. Harding, Joseph H. Marcus, Jared Carlson-Stevermer, Benjamin Kerr

## Abstract

**Abstract:** In the context of Wright’s adaptive landscape, genetic epistasis can yield a multipeaked or “rugged” topography. In an unstructured population, a lineage with selective access to multiple peaks is expected to rapidly fix on one, which may not be the highest peak. Contrarily, beneficial mutations in a population with spatially restricted migration take longer to fix, allowing distant parts of the population to explore the landscape semi-independently. Such a population can simultaneous discover multiple peaks and the genotype at the highest discovered peak is expected to fix eventually. Thus, structured populations sacrifice initial speed of adaptation for breadth of search. As in the Tortoise-Hare fable, the structured population (Tortoise) starts relatively slow, but eventually surpasses the unstructured population (Hare) in average fitness. In contrast, on single-peak landscapes (e.g., systems lacking epistasis), all uphill paths converge. Given such “smooth” topography, breadth of search is devalued, and a structured population only lags behind an unstructured population in average fitness (ultimately converging). Thus, the Tortoise-Hare pattern is an indicator of ruggedness. After verifying these predictions in simulated populations where ruggedness is manipulable, we then explore average fitness in metapopulations of *Escherichia coli*. Consistent with a rugged landscape topography, we find a Tortoise-Hare pattern. Further, we find that structured populations accumulate more mutations, suggesting that distant peaks are higher. This approach can be used to unveil landscape topography in other systems, and we discuss its application for antibiotic resistance, engineering problems, and elements of Wright’s Shifting Balance Process.

**Significance Statement:** Adaptive landscapes are a way of describing how mutations interact with each other to produce fitness. If an adaptive landscape is rugged, organisms achieve higher fitness with more difficulty because the mutations to reach high fitness genotypes may not be always beneficial. By evolving populations of *Escherichia coli* with different degrees of spatial structure, we identified a Tortoise-Hare pattern, where structured populations were initially slower, but overtook less structured populations in mean fitness. These results, combined with genetic sequencing and computational simulation, indicate this bacterial adaptive landscape is rugged. Our findings address one of the most enduring questions in evolutionary biology, in addition to, providing insight into how evolution may influence medicine and engineering.

## Introduction

The adaptive landscape was introduced by Sewall Wright to visualize potential constraints faced by evolving systems of genes (1). One incarnation of Wright’s landscape portrays the relationship between an organism’s genotype and its fitness as a topographical map. Imagine placing all possible genotypes of an organism together on a plane, where the distance between two genotypes represents the number of mutations needed to generate one genotype from the other (here we focus on asexual organisms). Each genotype is assigned a height directly proportional to its fitness (the third dimension). An evolving population is represented as a cloud of points on the resulting landscape, where each member of the population is a point. Novel genotypes arise in the population via mutations, expanding the extent of the cloud. In contrast, natural selection diminishes the range of the cloud, shifting its weight uphill as less fit genotypes are culled. Thus the combination of mutation and selection leads to the population “climbing” adaptive hills to their “peak,” which is a genotype from which all mutations are deleterious. If we assume strong selection and weak mutation (*SSWM*), the population cloud is mostly confined to a single climbing point, where the rapid fixation of each rare beneficial mutation shifts the point uphill (2, 3). Overall, a population’s evolutionary trajectory is taken to be sensitive to the gradients on this three-dimensional landscape.

As pointed out by many authors, including Wright (4–6), the actual geometry of the space of possible genotypes has extremely high dimensionality, which cannot be projected into two dimensional space in a way that preserves all distances. We explore an alteration of the classic representation (see (7–9)) that ensures genotypes differing by a single mutation are equidistant (while the distance between genotypes differing by multiple mutations is distorted). This approach involves creating a network, in which nodes are genotypes and edges connect genotypes differing by a single substitution. This network is embedded in two dimensions, where genotypes are grouped along the abscissa by their distance from a common genotype and along the ordinate by their fitness.

Using this representation, a mutation network without epistasis is shown in Figure 1a. This network would also be labeled as a “smooth landscape,” as the single peak is accessible (i.e., can be reached by a series of beneficial mutations) from any other genotype. Evolution of populations on this landscape gives an example of mutational convergence. Under *SSWM* assumptions, we show that despite three different initial mutational steps, independent evolutionary trajectories converge at the peak (Fig. 1a) and fitness likewise converges (Fig. 1b). In contrast, Figure 1c shows a network with sign epistasis, where the sign of the fitness effect of a mutation depends on the background in which it occurs. Such sign epistasis is a necessary (but not sufficient) condition for the existence of multiple peaks. On this “rugged” landscape, the final genotype reached under three independent trajectories is contingent upon the initial mutation (Fig. 1c). In this case, fitness values of different populations can remain divergent over time if peaks are heterogeneous in height (Fig. 1d). Here we see that a population can become trapped at a suboptimal peak in the presence of epistasis.

**Figure 1.**
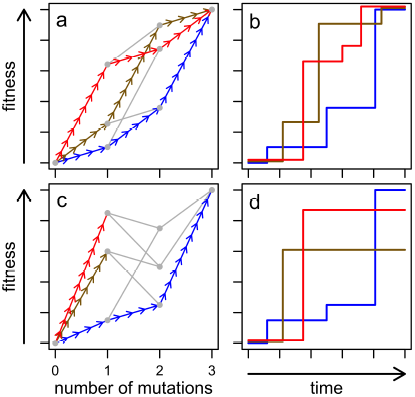
**Adaptive paths in hypothetical landscapes**. Here we consider a simple biallelic three locus system. (a) The adaptive landscape can be visualized by plotting genotype fitness as a function of the number of mutations on a wild type background. Each of the 2^3^=8 genotypes is given by a gray point and edges (arrows or gray lines) connect genotypes differing by a single mutation. An adaptive peak is a genotype from which all mutations are detrimental. A hypothetical landscape with a single peak (the triple mutant) is shown here. A selectively accessible path exists between two genotypes if a series of beneficial mutations connects the less fit genotype to the more fit genotype. On this “smooth” landscape, all of the 3!=6 paths between the wild type (lacking mutations) and the triple mutant are selectively accessible; three of these paths are shown by the arrows in different colors. (b) Average fitness over time is shown for three possible populations following the paths in part a. If we assume that selection is strong and mutation is weak, we can represent the fixation of each beneficial mutation as a step up in the fitness trajectory. All trajectories converge on the same final fitness value. (c) A hypothetical landscape with multiple peaks. Starting with the wild type, selection can take the population to different adaptive peaks on this “rugged” landscape, as illustrated by the different colored trajectories. (d) Average fitness over time is shown again for three possible populations following the paths in part c. The final fitness of different evolving populations can vary.

Because Sewall Wright thought epistasis was pervasive (10), he was particularly concerned about confinement of populations at suboptimal peaks within rugged landscapes. He proposed the Shifting Balance Process (SBP) to explain how populations move from lower to higher peaks. Integral to the SBP is population structure. Wright envisioned a population that was distributed into semi-isolated, sparsely-populated subpopulations (demes) in which genetic drift enabled some subpopulations to take downward steps by fixing deleterious mutations. In this way, a subset of the metapopulation is able to move from one peak’s domain of attraction to another, thus crossing “adaptive valleys.” Therefore, Wright’s SBP depends on two critical assumptions: the presence of epistasis generating landscape ruggedness and the presence of population structure.

By using one factor discussed by Wright (population structure) as an experimental variable, an empirical assay can be constructed for another of Wright’s factors (landscape ruggedness). Upon first glance, population structure would seem to hinder adaptation. In a population in which migration is not spatially restricted (unstructured population), a beneficial mutant that arises can rapidly fix in what is termed a selective sweep. On the other hand, a favored mutant arising in a population with restricted migration (structured population) advances more slowly in what might be termed a “selective creep.” By reducing the rate of initial adaptation, the slow competitive displacement occurring within a structured population may also allow multiple semi-independent searches of the fitness landscape by geographically distant regions of the population. For a smooth landscape (e.g., Fig 1a), this enhanced exploration is superfluous as all selectively accessible trajectories lead to the same single peak. Therefore, on smooth landscapes, structure only slows adaptation. However, on a rugged landscape, additional exploration may reveal alternate peaks. For instance, in Fig. 1c, while an unstructured population might exclusively follow one of the colored trajectories, a structured population may be able to explore them all simultaneously. In this way, a structured population can survey a broader set of paths. As discovered peaks may differ in height, a comparison of them enables the population to eventually reach a better endpoint on average (11, 12). On a rugged landscape, fitnesses in populations differing in structure emulate the classic Tortoise-Hare fable. Specifically the unstructured population initially adapts faster (the Hare) but is overtaken by the structured population (the Tortoise), which is a poor starter but a strong finisher. Importantly, on a smooth landscape, the Tortoise never takes the lead, and the crossing of average fitness trajectories is not predicted. Thus, when manipulations to population structure do produce a Tortoise-Hare pattern, we have a signature of ruggedness.

Before investigating this signature in a biological system in which landscape topography is cryptic, we confirm the above predictions using a computational system in which landscape topography is known and manipulable. Specifically, we control landscape ruggedness by employing Kaufmann’s NK model (13–15) and then track evolving metapopulations of bit strings, in which the pattern of migration between demes is manipulated. Following this simulation, we then turn to evolving metapopulations of *Escherichia coli* under a similar experimental manipulation of population structure. Discovery of a Tortoise-Hare pattern would be indicative of a rugged topography.

## Results and Discussion

### Patterns of Average Fitness

In the NK model, simulated organisms are bit strings of length N, and the parameter K is the number of loci affecting the fitness contribution of each locus (see Methods). As K increases, the level of epistatic interaction increases, yielding more rugged landscapes; hereafter, we refer to K as a “ruggedness” parameter. We explore how ruggedness affects fitness trajectories in evolving metapopulations that differ in population structure. We consider either metapopulations with spatial restrictions to migration (hereafter, the Restricted treatment) or metapopulations where migration can occur between any two demes (Unrestricted treatment). For a smooth landscape topography (K=0, N=15), average fitness initially increases more rapidly in the Unrestricted treatment relative to the Restricted treatment; however, both trajectories converge over time (Fig. 2a). For a rugged landscape (K=8, N=15), fitness in the Restricted treatment once again lags behind fitness in the Unrestricted treatment at the outset. Instead of converging, however, the fitness trajectories cross, yielding a higher final fitness for the spatially restricted treatment (Fig. 2b). Indeed, we find significantly higher fitness in the Restricted treatment for K>3 at the end of our simulation (Fig. 2c; Mann-Whitney tests with Bonferroni corrections, p<0.001). The pattern in Figure 2b agrees with the Tortoise-Hare prediction, while the crossing does not occur for the Unrestricted treatment in Figure 2a. More generally, with sufficient ruggedness, a structured population can eventually outperform an unstructured population (Fig. 2c). This pattern is not limited to our specific form of population structure, as Bergman *et al*. (1995) (16) reported a similar result using an NK model where bit strings dispersed variable distances along a single dimension.

**Figure 2.**
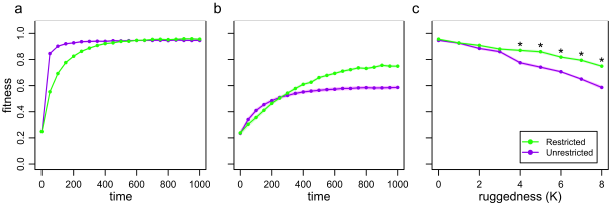
**Fitness in the NK model**. Metapopulations of bit strings of length N=15 evolved where either migration was restricted to occur between neighboring demes or migration was unrestricted (occurring between any two demes). (a) Average fitness in the metapopulation is shown over time on a smooth landscape (K=0) or (b) a rugged landscapes (K=8). (c) Average fitness at time point 1000 is shown as a function of the ruggedness parameter, K. Note the values at K=0 and K=8 in part c correspond to the values at time point 1000 in parts a and b, respectively. In all plots, points represent the mean of 50 replicates, shading gives the standard error of the mean and asterisks indicate significant differences.

We next turned to examining fitness trajectories in evolving metapopulations of *Escherichia coli*. Similar to the NK model, we propagated the bacteria under two treatments differing in migration pattern: Restricted and Unrestricted (see Methods). Early and late in the evolutionary run, we sampled five random isolates from each metapopulation and determined their fitness relative to the common ancestor. Early in the experiment (at transfer 12), fitness in the Restricted treatment was significantly lower than the Unrestricted treatment (Fig. 3; Mann-Whitney test, p=0.015). However, at the end of the experiment (transfer 36), the opposite pattern was found, with fitness in the Restricted treatment surpassing the Unrestricted treatment (Fig. 3; Mann-Whitney test, p=0.015). This pattern is consistent with a rugged landscape topography.

**Figure 3.**
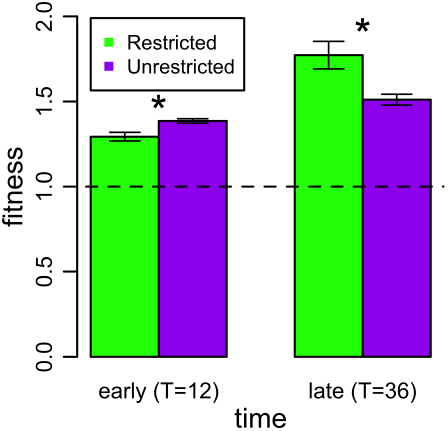
**Bacterial fitness**. Metapopulations of bacteria evolved where migration was spatially restricted or unrestricted. Average relative fitness of five isolates randomly sampled from the metapopulation is shown early in the experiment (at transfer T=12) and late in the experiment (at transfer T=36). As in Figure 2b, the ordering of fitnesses for the two treatments flips over time. Bars represent the mean of 5 replicate metapopulations, whiskers give the standard error, and asterisks denote significant differences.

### Patterns of Evolutionary Distance

There are a few ways to account for the benefit that population structure confers on rugged landscapes (Figs. 2b and 3). First, a population may have access to multiple peaks that differ in height. A structured population can explore multiple domains in parallel, eventually comparing the results. Thus, it will tend to attain a higher endpoint; for the same reasons that the expectation of the maximum of a sample increases with sample size. This effect holds when all peaks are equidistant from the ancestral population. A second possibility (not mutually exclusive with the first) is that peaks differ in both height and distance from the ancestral population. Suppose that the initial mutations on accessible paths to the more distant and higher peaks are less beneficial than mutations leading to the nearby peaks, as in (17). In this case, intermediate genotypes approaching distant peaks risk being outcompeted in an unstructured population (consider the first mutant on the blue path to the more distant peak in Fig. 1c competing against the other first mutants). This is because the slower fixation of these intermediates allows for better competitors (from domains of nearer peaks) to arise. In contrast, these more distant peaks become accessible in a structured population due to reduced competitive displacement. If some of these distant peaks are also higher, then structured populations are predicted to both achieve better fitness and accumulate more mutations.

To explore the number of mutations accrued by evolving populations, we first return to the NK model. We define evolutionary distance to be the number of mutational differences between an evolved isolate and its ancestor. In the NK model, this is the Hamming distance (18). As ruggedness increases, the degree of population structure affects final evolutionary distance from the ancestor; we find a significantly higher distance in the Restricted treatment for K>3 at the end of our simulation (Fig. 4; Mann-Whitney tests with Bonferroni corrections, p<0.001). Thus, on a rugged landscape, a population with restricted migration moves both higher (Fig. 2c) and further (Fig. 4) than a less structured population.

**Figure 4.**
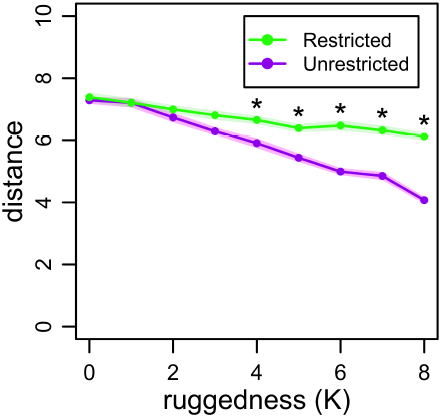
**Distance in the NK model**. Metapopulations of bit strings of length N=15 evolved where migration was spatially restricted or unrestricted. Evolutionary distance is the number of bits differing between an evolved isolate and its ancestor (the Hamming distance). Average distance at time point 1000 is shown as a function of the ruggedness parameter, K. Points represent the mean of 50 replicates, shading gives the standard error of the mean, and asterisks denote significant differences.

Evolutionary distance was explored similarly in the *E. coli* metapopulations. In addition to assessing fitness on the isolates from the end of the experiment (transfer 36), we sequenced their genomes to determine their evolutionary distance from the common ancestor. The locations of all identified mutations in all sequenced isolates are shown in Figure 5. The number of mutations accumulated in each isolate is the evolutionary distance (the points to the right of the table in Fig. 5). We find that isolates have moved a significantly greater distance in the Restricted treatment (Mann-Whitney test, p=0.045). The increased distance traversed by digital and bacterial populations is consistent with rugged landscapes in which more distant peaks are being reached by structured populations.

**Figure 5.**
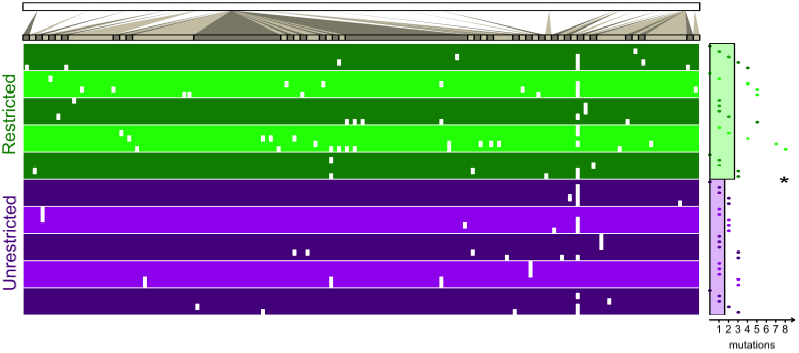
Bacterial distance. Metapopulations of bacteria evolved where migration was spatially restricted or unrestricted. At the end of the experiment, five isolates from each metapopulation were sequenced at the genome level. The top bar represents the genome of *Escherichia coli*. Genome regions with mutations are magnified for the table. Each isolate is a single row in the table and the location of each of its mutations is indicated by a white mark. The five isolates from each of the five replicate metapopulations are grouped by alternating shades of green (for the Restricted treatment) or purple (for the Unrestricted treatment). The horizontal distance of the point to the right of the table denotes the number of mutations in the isolate (its evolutionary distance). The horizontal distance of the green bar and the purple bar to the right of the table gives the average distance (the average of replicate averages) of isolates from the Restricted and Unrestricted treatments, respectively. The asterisk denotes a significant difference between treatments. More information regarding each mutation can be found in the Supplement.

The ability of the structured populations to move higher (in average fitness) and further (in evolutionary distance) is engendered by the capacity for parallel search. The presence of simultaneous selective creeps should increase the standing diversity within a structured population relative to an unstructured one. In line with this prediction, the metapopulations in the Restricted treatment had significantly higher genotypic diversity than the Unrestricted treatment (Mann-Whitney test on the nucleotide diversity index **p**, p=0.016). We note that greater diversity in structured populations is expected regardless of the topography of the landscape (see Supplemental Figure 1), however, such diversity is only advantageous when the landscape is multi-peaked.

### Previous Empirical Work

An extreme form of population structure involves a set of completely isolated populations. If the landscape contains multiple peaks of different heights, these populations can diverge in fitness, genotype and phenotype. Korona *et al*. (19) and Melnyk & Kassen (20) measured phenotypic diversity within, and diversity in fitness among, a set of replicate evolving microbial populations. Under some growth conditions (growth on agar surfaces for *Rastonia eutropha* (19) and growth in minimal xylose for *Pseudomonas fluorescens* (20)) these authors found that both forms of diversity remained high at the end of the experiment, which they took as evidence for a rugged landscape. Under other growth conditions (growth in liquid for *R. eutropha* (19) and growth in minimal glucose for *P. fluorescens* (20)) they found lower diversity, consistent with a smoother landscape. Like the present study, these earlier studies used statistical patterns to infer topographical properties of landscapes.

This statistical approach was also used in a recent study by Kryazhimskiy *et al*. (21). They propagated metapopulations of asexual *Saccharomyces cerevisiae* and varied the rate (as opposed to pattern) of migration. In contrast to the results we report, they find yeast from treatments with higher rates of migration evolved higher fitness over the course of their experiment. This led the authors to conclude that epistasis was weak and the landscape was smooth. While it is entirely possible that their yeast and our bacteria differ in landscape topography, we discuss some alternative interpretations of their results in the Supplement.

Rather than inferring landscape topography from population-level statistical patterns, an alternative approach involves fully characterizing the fitnesses of genotypes in a small section of the landscape (22–30). This approach often involves considering two genotypes differing by *M* mutations, engineering all 2*^M^* possible combinations, and assessing the fitness of each constructed genotype. Epistasis can be gauged directly by measuring the influence of genetic background on the fitness effect of a mutation. Some studies using this approach have uncovered instances of sign epistasis (22, 24, 29), while other studies have found only magnitude epistasis (23, 26). Some studies have reported multiple peaks (24, 29), while others have found only a single peak (22, 26). As with the inferential approach described above, this engineering approach has revealed a potential diversity of landscape topography. We suggest that combining the (top-down) inferential approach and (bottom-up) engineering approach is a promising direction for exploring the nature of adaptation.

### Wright’s Shifting Balance Process

During the Modern Synthesis, two divergent views on adaptation were debated. One view considers adaptation as the sequential fixation of beneficial mutations. This perspective (often associated with Fisher) does not highlight epistatic contingency, focusing instead on selection and mutation as the major processes of evolution (31). A second perspective (often linked to Wright) recognizes epistatic interaction as constraining adaptation. Based on his empirical observations, Wright felt that such epistasis was pervasive, leading, within his adaptive landscape metaphor, to multiple peaks (2). In this view, evolving populations can become trapped in suboptimal positions. To illustrate how escape from the domains of suboptimal peaks was possible, Wright introduced the Shifting Balance Process (SBP). He assumed that adapting populations were spatially structured as a metapopulation of semi-isolated demes. The SBP is heuristically divided into three phases. In Phase I, the demes, which Wright assumed were small, drift in genotype space, enabling movement into other domains. In Phase II, selection within demes produces a set of semi-independent hill-climbing episodes. In Phase III, the resulting genotypes are exchanged among demes and the best competitors spread through the metapopulation at large. Thus, Wright’s view of adaptation, in contrast to Fisher’s, invokes a complex combination of processes, specifically brought together to solve a problem generated by epistatic contingency.

While there have been various theoretical explorations of the plausibility of SBP (16, 32–35), Wright’s ideas have been criticized because they demand a delicate balance of various evolutionary processes. For instance, populations need be small enough for effective drift (Phase I), but large enough for effective selection within demes (Phase II). Migration should be sufficiently restricted for drift and selection within demes (Phase I and II), but sufficiently unrestricted for effective exchange of genotypes among demes (Phase III). However, if we abandon Wright’s goal of explaining how populations cross valleys, many of these conflicts vanish. Both Fisher and Wright acknowledged that environmental change could alter the landscape, and, in the process, reposition the peaks. Imagine that a population experiences such a change and subsequently resides somewhere on the new landscape with access to multiple domains. In our experiment, our ancestor contained deleterious mutations and evolved in a stressful environment (see Methods), which potentially yielded access to multiple peaks. In this case, demes need not be small for the discovery of multiple peaks (and indeed, our experimental demes were large). With large subpopulations, selection within demes will proceed efficiently; however, limitations to migration between demes will still allow for parallel exploration of a rugged landscape. Thus, Phases II and III can jointly yield adapted populations even if Phase I is absent. If landscapes are indeed rugged, population structure can retain the critical role Wright foresaw, even if all the details of the SBP are not present.

### Applications

One case where populations are potentially poised in multiple domains on a landscape involves the evolution of microbes exposed to antibiotics. When a bacterial population experiences a sufficiently high concentration of an antibiotic, susceptible genotypes are replaced by resistant mutants. When the drug is removed, these mutants tend to carry fitness costs relative to their susceptible progenitors.

The cost can be alleviated by a mutation resulting in reversion to susceptibility or a mutation that compensates for the impairment without loss of resistance (36, 37). There is some evidence that reversion and compensation constitute distinct peaks in a rugged landscape (38, 39). Thus, we see that a changing environment (exposure and removal of a drug) may position a microbial population at a landscape position where multiple peaks are accessible (7). It is at such a position that population structure may influence the evolutionary trajectory. Björkman *et al*. (38) and Nagaev *et al*. (39) serially passaged *Salmonella typhimurium* and *Staphylococcus aureus* resistant to fusidic acid either in well-mixed flasks or within murids (mice or rats). These authors found that the bacteria more often reverted when grown *in vivo* than *in vitro*. They explain these results by noting that the flask and murid environments differ markedly and may consequently place different selective pressures on revertants and compensated strains (indeed, they present data to this effect). In our terminology, the landscape in a flask and a mouse may be different. However, even if the landscape was identical (but rugged) in both, the results might not be unexpected because a murid environment is highly structured and a shaking flask is not. Thus, if the “reversion peak” is higher than most to all of the “compensation peaks” (the authors present data consistent with this ordering) then evolution in a structured environment is predicted to revert at higher frequency. In this way, the structure that pathogenic bacteria experience (including in the bodies of human hosts) can potentially influence the course of antibiotic resistance evolution.

Not only can the ideas in this paper apply in a medical context, but also they may address practical engineering problems. Evolutionary principles have been utilized to find solutions to computational problems, a discipline known as evolutionary computation. In this field, putative solutions constitute a population, new solutions are generated by mutation and recombination, and better solutions can outcompete their contemporaries. One defining feature of a difficult problem is the presence of multiple optima in the map from the specification of a solution (i.e., genotype) to its quality (i.e., fitness). As early as 1967, Bossert (40) suggested that dividing a population of solutions into subpopulations could yield better evolutionary outcomes. Subsequently, the inclusion of population subdivision in evolutionary algorithms has produced better solutions in a variety of applications, including analogue circuit design (41), financial trading models (42), and multi-objective scheduling (43). Besides the efficiency in networked computational resources that accompanies population subdivision, a deeper exploration of the landscape of solutions is predicted to occur when multiple domains can be semi-independently searched (44, 45).

### Synthesis

Contingency in evolution is affected by the underlying topography of adaptive landscapes. When no epistasis is present, a smooth landscape results, and ultimately evolution converges to the single peak. Contingency requires (sign) epistasis, specifically of a kind generating multi-peaked, or rugged, landscapes. (We again note that sign epistasis is not sufficient for multi-peaked landscapes and indeed can constrain evolution to a subset of paths to a single peak; see Weinreich *et al*. (22) for an example.) In the case of heterogeneous peak heights, population structure may enable the simultaneous exploration of multiple domains and ultimately lead to the discovery of higher peaks than would be possible in an unstructured population. Thus, using population structure as an experimental variable, we have a signal for this kind of ruggedness. In the case of our bacterial populations, we have presented a pattern consistent with ruggedness. Additionally, it appears that structured populations move further in the landscape, suggesting that the most accessible peaks may not be the highest. Ultimately it is an empirical issue whether other biological systems posses such ruggedness. However, statistical patterns in fitness and evolutionary distance may help to distinguish landscape topography. Specifically, a rugged landscape topography can be inferred by comparing structured (Tortoise) and unstructured (Hare) populations and assessing whether slow and steady adaption “wins the race.”

## Methods

### Ancestral strain

The bacterial ancestor was derived from a K-12 strain of *E. coli* (BZB1011) by selecting for resistance to colicin E2, then colicin D, and then phage T6. Both resistance to colicin E2 and phage T6 are known to be individually costly (46, 47). The initiation of the experiment with an unfit strain was intentional (see Supplement for rationale).

### Experimental treatments

Each metapopulation was comprised of 96 subpopulations (the 96 wells of an 8×12 microtiter plate). The metapopulation was initiated with the ancestral strain in each well. These subpopulations grew for 12 hours in 200 mL of lysogeny broth (LB- Miller) supplemented with a sub-inhibitory concentration of tetracycline (0.25 mg/mL). After growth, each well in the metapopulation was diluted 40-fold into fresh growth medium using a 96 slot-pin multi-blot replicator (5 µL into 200 µL). Immediately following dilution, migrations among wells occurred. Migration was either restricted to occur between subpopulations adjacent to each other or was unrestricted. In both treatments, every well had 1/3 probability of experiencing an immigration event from one random well in its neighborhood. In the Restricted treatment, this neighborhood included the wells directly north, east, south or west of the focal well (using periodic boundary conditions to eliminate edge-effects). In the Unrestricted treatment, the neighborhood included all wells minus the focal well. All migration events were executed by a BioRobot 8000 liquid-handling robot (Qiagen), which transferred 5 µL from the source well within the plate from the previous transfer into the destination well within the plate from the current transfer. Between transfers, plates were incubated (37 °C) and shaken (350 rpm using a microtiter shaker). Each metapopulation was propagated for a total of 36 transfers and each treatment contained five replicates.

### Competition assay

We chose five random isolates from the last transfer of each metapopulation (here we denote any one of these strains as *E*). We marked our ancestor with resistance to phage T5 (we denote this marked ancestor as *A*). Before the competition, *E* and *A* were grown separately in 200 µL of growth medium for two 12-hour cycles (with 40-fold dilution at transfer). After this acclimation phase, we added 5 µL of *E* and 5 µL of *A* to a well containing 200 µL of fresh growth medium. The titer of each strain was assessed, through selective plating with and without phage T5, immediately after the competition was initiated and again after 12 hours. If *X*(*t*) is the titer of strain *X* at time *t*, then the fitness of the evolved strain relative to its ancestor is given by:

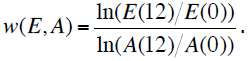

### Whole Genome Resequencing

Using the same isolates from the last transfer, we performed chromosomal DNA extractions using Qiagen Mini DNA Kits. Each sample was barcoded and multiplexed to 24 samples per lane with Illumina TruSeq. Whole genome resequencing (University of Washington High Throughput Genomics Unit) was performed with single-end 36-bp unpaired reads on Illumina HiSeq to an average of 30X coverage. Illumina reads were aligned for mutational discovery by Breseq 0.19 (J. E. Barrick, unpublished algorithm) against *E*. *coli W311* [GenBank: AP009048]. Alignments were considered only if they covered 95% of a read. For every isolate, Sanger sequencing (Genewiz) of several loci (*fimE*, *marR*, *ompF*, and *stfR*) was used to confirm putative mutations.

### NK Model

For the simulations, individuals were embedded within 96 demes in an 8 x 12 array. Each deme contained 1000 organisms. Each organism’s genotype was a bit string (fixed length ordered list of 0’s and 1’s) of length N=15. The fitness of an organism was the sum of the fitness contributions of each of the 15 loci divided by the number of loci. The contribution of each locus was determined by its allelic state and the allelic states of the subsequent K loci (wrapping to the beginning of the bit string as needed). For each locus, 2^K+1^ random numbers (uniformly distributed between 0 and 1) described all possible fitness contributions of that locus (given any possible combination of alleles at relevant loci). Thus a mutation at a single locus affected the fitness contribution of the mutated locus and K other loci. Selection within a deme involved the removal of a random organism, regardless of fitness, and its replacement by the birth of an organism from the same deme chosen by a fitness-weighted lottery. Upon birth, the offspring bit string differed from its parent at a random locus with probability 0.1 (the mutation rate). This Moran death-birth process was iterated 1000 times for each deme, followed by migration between demes. During each migration event, 25 individuals were chosen at random and removed from one deme (the destination), and then replaced by copies of 25 individuals chosen at random from the other deme (the source). Each deme experienced an immigration event with probability 1/3. Migration was either restricted or unrestricted in precisely the same manner as the bacterial experiment above. Each replicate run of the Unrestricted treatment was paired with a replicate of the Restricted treatment, where each member of the pair shared the same NK landscape as well as the same ancestor (a random bit string used to populate the entire metapopulation). In the figures, one selection-migration episode is termed an “update.”

## Acknowledgments

We thank A. Covert and C. Wilke for sharing ideas that inspired this work; and P. Conlin, B. Connelly, J. Cooper, K. Dickinson, C. Glenney, S. Hammarlund, H. Jordt, K. Raay, and S. Singhal for comments on the manuscript. This material is based in part on work supported by the National Science Foundation under Cooperative Agreement Number DBI-0939454, a National Science Foundation Graduate Research Fellowship (to J.R.N.), a NSF CAREER Award Grant (DEB0952825), and a UW Royalty Research Fund Award (A74107).

## Supplementary Information

### Diversity in digital and bacterial populations

A structured population performs a broader search on the adaptive landscape, as the rate of competitive displacement is lower. Consequently, the standing genetic diversity of a structured population is expected to be greater than diversity in an unstructured population. In Supplemental Figure 1a, we see that this pattern does not depend on landscape ruggedness (Mann-Whitney tests, p<0.01 for K=0 and K=8). Thus, despite landscape topography, we predict to find higher genetic diversity in a structured population, and this is what we find in our bacterial metapopulations (Mann-Whitney test, p=0.015; Supp. Fig. 1b).

### Diversity methods

Consider a sample of *G* genotypes (bit strings or nucleotide sequences). We use the diversity index of Nei and Li (1979) (1):

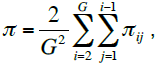

where π*_ij_* is average number of differences (in bits or bases) per site between genotype *i* and genotype *j*. We refer to π as bit diversity (in the NK model) or nucleotide diversity (for our bacterial system).

**Figure S1:**
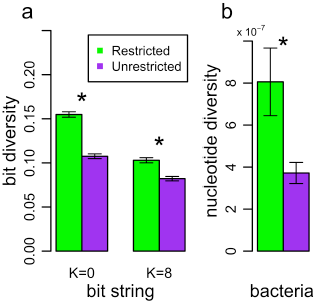
Genetic diversity in the digital and bacterial populations. (a) Average bit diversity of a sample of eight evolved bit strings from time point 1000 in the NK model simulations. Whether the landscape is smooth (K=0) or rugged (K=8), diversity is significantly greater in the Restricted treatment than the Unrestricted treatment. Bars represent the mean of 40 replicates. (b) Average nucleotide diversity within bacterial metapopulations at the final transfer of the experiment (T=36). For each metapopulation, full genome sequences from each of five isolates was used to compute the diversity index. Nucleotide diversity is significantly greater in the Restricted treatment than the Unrestricted treatment. Bars represent the mean of 5 replicates. In both parts of the figure, whiskers give the standard error and asterisks indicate significant differences.

### How a rugged landscape can fail to give a Tortoise-Hare signal

On a rugged landscape, fitness in a structured population will increase more slowly than an unstructured population (the Tortoise initially lags behind, before overtaking, the Hare; see Fig. 2b). That is, for populations differing in structure evolving on a rugged landscape, early evolution will produce a pattern similar to that predicted under a smooth landscape (e.g., before time point 250, the pattern in Figure 2b would be hard to distinguish from the entire trajectory of Figure 2a). While the presence of a Tortoise-Hare pattern indicates ruggedness, its absence does not necessarily imply a smooth landscape.

For example, in the experiment of Kryazhimskiy *et al*. (2012) (2), the unstructured population ended the experiment with higher average fitness. This pattern is consistent with a smooth landscape, but is not inconsistent with a rugged one. As the authors themselves acknowledge, had their experiment run longer, they may have observed higher fitness under lower rates of migration (i.e., a fitness crossing).

Even when there is abundant time for evolution to take place, it is still possible that evolution on a rugged landscape will fail to yield the Tortoise-Hare pattern.

For instance, it is possible that the landscape is rugged, but peaks are of a homogeneous height. This could produce the fitness pattern shown in Figure 2a.

Finally, the landscape could be rugged with heterogeneous peak heights, but the ancestor could be positioned in the domain of a single peak. Consistent with this possibility, Kryazhimskiy *et al*. started their experiment with a lab-adapted strain of yeast and evolved their populations under standard laboratory conditions. If their yeast had access to only a single domain in a rugged landscape, a Tortoise-Hare pattern would not be expected. In such a case Kryazhimskiy *et al*. would be justified in claiming that the *local* topography of such a landscape was smooth (and indeed, they restrict their claim of smoothness accordingly).

As outlined in our Methods, we introduced several deleterious mutations into our ancestor and evolved our populations under a stressful environment (in the presence of sub-lethal concentrations of the antibiotic tetracycline). Such manipulation was intended to displace our ancestral genotype from a peak, but it also may have placed it at a point where multiple domains were accessible. To address the effect of ancestor starting position, we describe additional NK simulations here. In addition to starting our ancestor at a random bit string, we consider three other starting positions: (i) valley, (ii) pre-adapted, and (iii) “silver-spoon.” For the valley simulations, we performed a “hill-plunge of steepest descent,” moving downhill from a random genotype until we hit a valley genotype (a genotype from which all mutations were beneficial), which served as the ancestor. For the pre-adapted simulations, we allowed a random ancestor to evolve briefly (in an unstructured population) to produce a “pre-adapted” ancestor. For the silver-spoon simulations, all genotypes were ranked for fitness and the genotype defining the 99^th^ fitness percentile was chosen as the ancestor.

In the random and valley starting positions, the Tortoise-Hare pattern was observed and the Structured treatment ended at significantly higher average fitness than the Unrestricted treatment (Mann-Whitney tests, p<0.001; Supp. Fig. 1b). However, in the pre-adapted and silver-spoon starting positions, the Tortoise-Hare pattern was not seen and fitness was indistinguishable between the treatments in the long run (Mann-Whitney test, p=0.36 and p=0.57 respectively). These simulations demonstrate that the starting position of a population in a landscape will influence the statistical pattern of fitness of populations differing in structure.

### Additional NK simulation methods

To study the effect of starting position in the adaptive trajectories in structured and unstructured populations, we examined starting the population with different types of ancestors. The “random” ancestor (used in the primary text) is simply a random bit string. To generate the “valley” ancestor, we start with a random bit string and substitutes the worst (lowest fitness) possible mutation until no deleterious mutations are possible, and that bit string is the ancestor for the evolutionary run. To produce the “pre-adapted” ancestor, we start with a random bit string and evolve a population initialized with this bit string under unrestricted migration for 50 updates. Then bit strings are sampled from the evolved population until one is found that has a higher fitness than the starting bit string. That adapted bit string is the ancestor. To determine the “silver-spoon” ancestor, all possible bit strings (2^15^) are ranked according to fitness and the 99^th^ percentile genotype is the ancestor.

**Figure S2:**
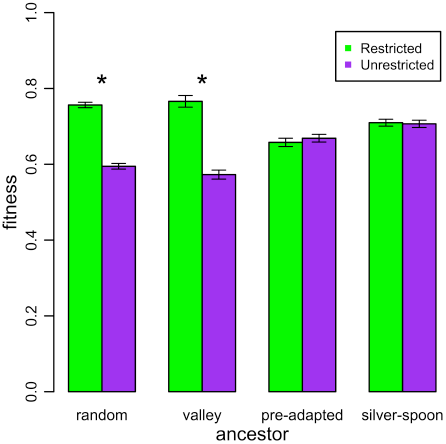
Metapopulations of bit strings of length N=15 evolved on a rugged landscape (K=8), where migration was either restricted or unrestricted. Average fitness in the metapopulation is shown at time point 1000 for a randomly chosen ancestor, an ancestor starting in a valley, an ancestor resulting from adaptation before the run, and an ancestor in the top percentile of fitness (the “silver spoon” ancestor). Bars represent the mean of 40 replicates, whiskers give the standard error of the mean, and asterisks indicate significant differences.

**Table S1:**
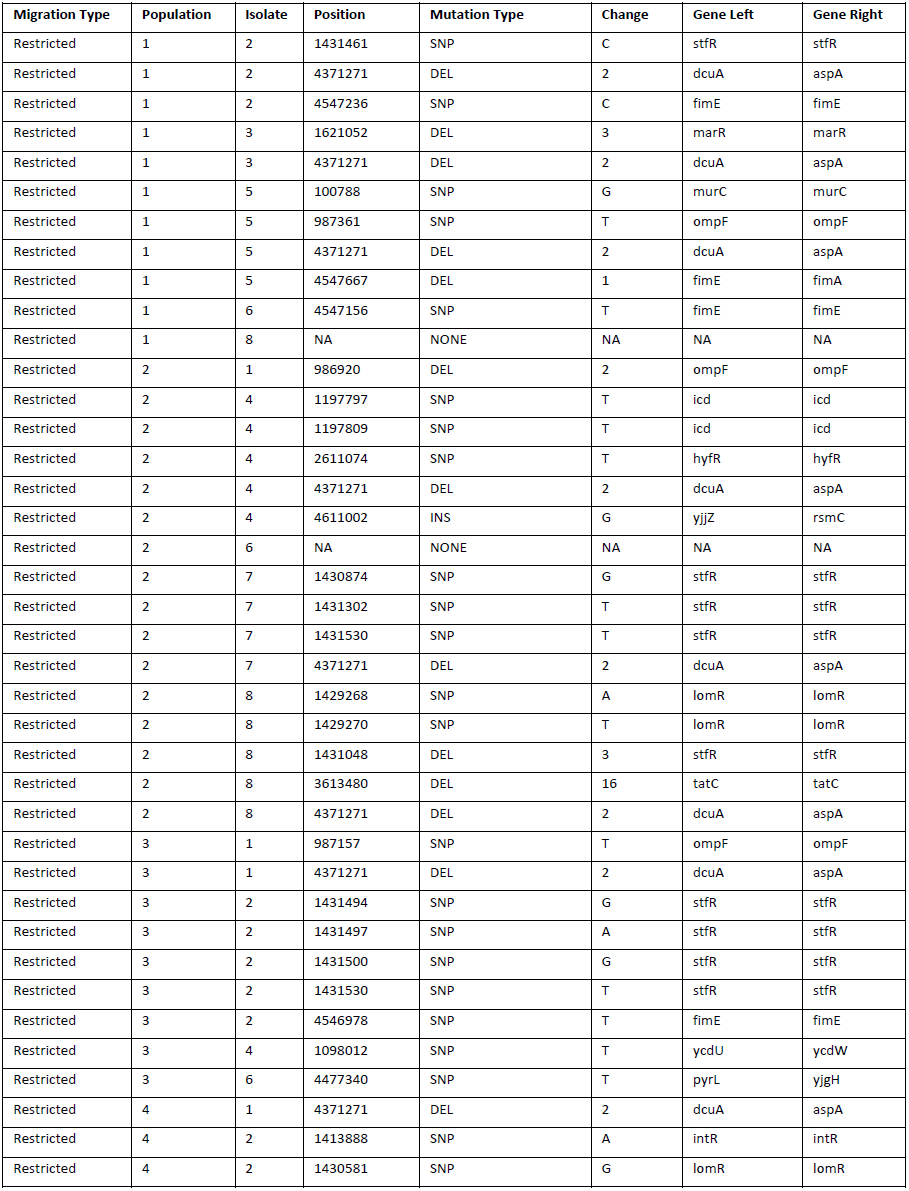
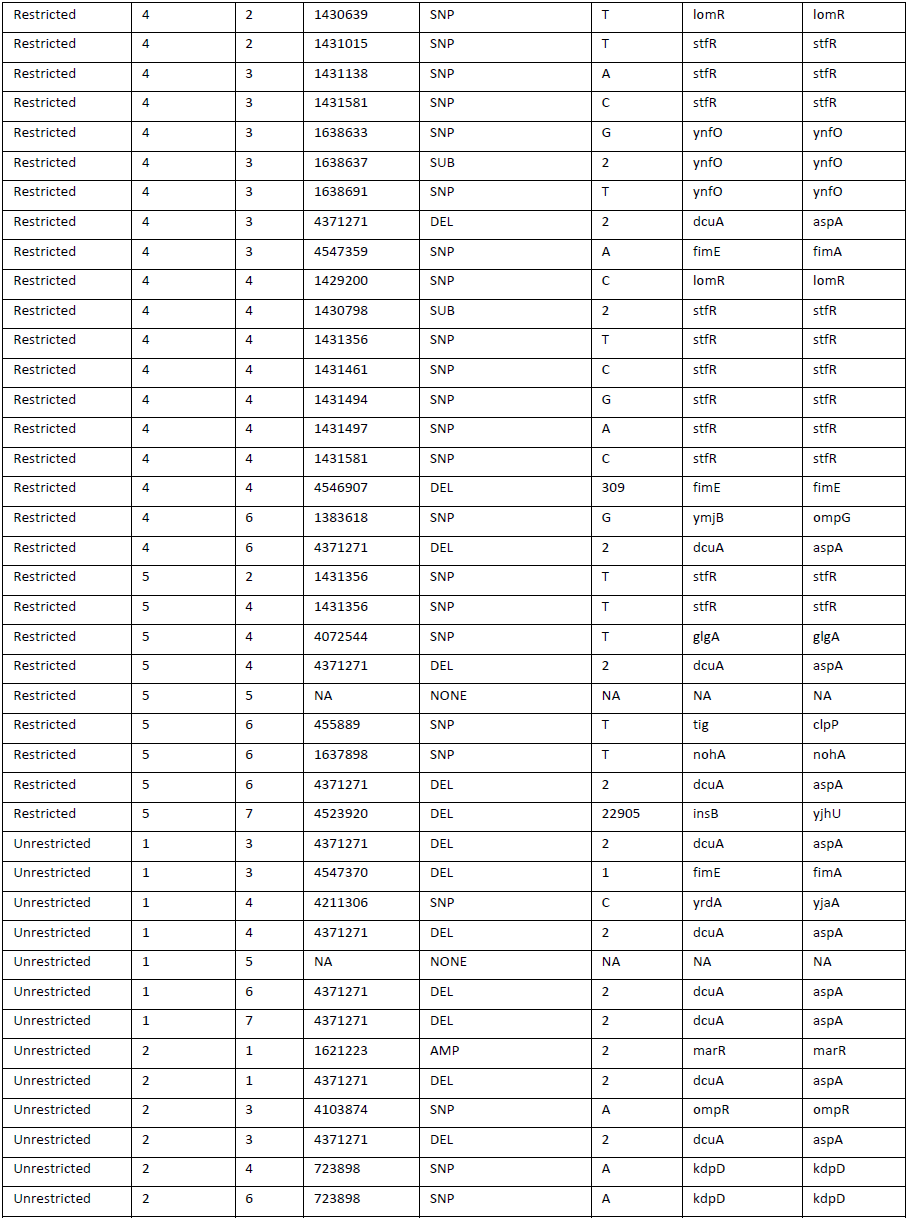
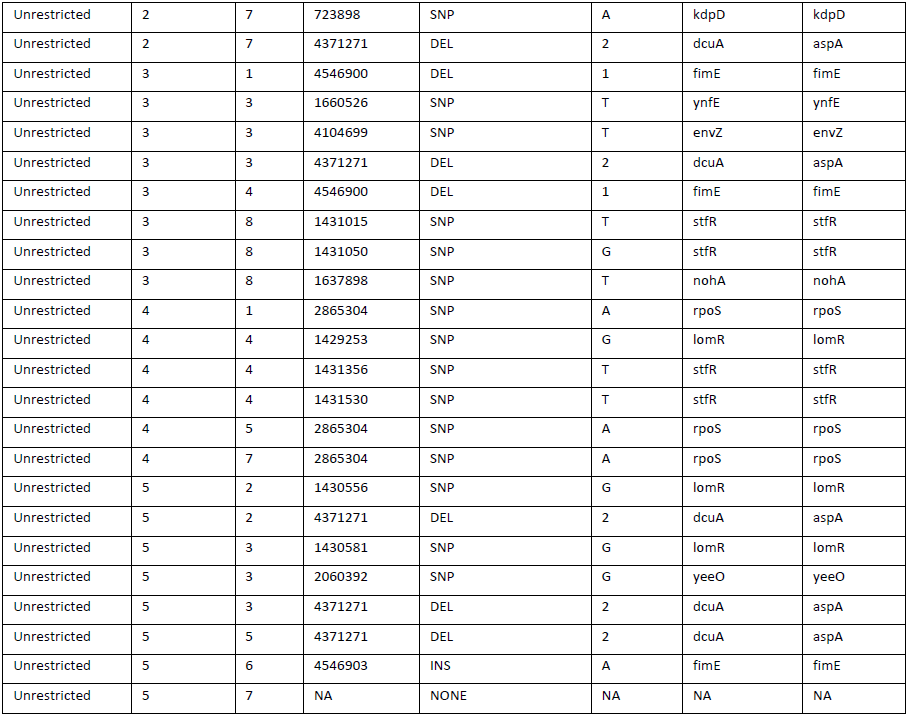

Each row denotes a mutation discovered in the evolved (transfer 36) isolates that was not present in the ancestor. The columns are as follows: Migration Type (the pattern of migration defining the treatment of the isolate), Population (the identifier for the replicate metapopulation (1–5) of the isolate), Isolate (the identifier for the mutation’s isolate (1–8)), Position (the genomic location of the mutation according to *E*. *coli W311* [GenBank: AP009048]), Mutation Type (Single Nucleotide Polymorphism (SNP), Deletion (DEL), Insertion (INS), Substitution (SUB) or Amplification (AMP)), Change (the length of a mutation for DEL, or AMP, or the new nucleotide state for SNP and INS), Gene Left (the nearest open reading frame to the prior to the mutation), Gene Right (the nearest open reading frame after the mutation). Note that if Gene Left and Gene Right are the same, the mutation falls within that gene.

### Violation of Strong-Selection-Weak-Mutation (SSWM) Assumptions

The relatively high degree of diversity in the evolving bacterial populations demonstrates a violation of strict SSWM assumptions. Isolates from the same metapopulation (in both treatments) were often not single mutant neighbors (see Figure 5 and Supplemental Table 1); thus, the diversity does not simply represent a mixture of the genotype fixing and the genotype being displaced in the midst of selective sweep. While divergent genotypes are expected in the structured population (Restricted Migration), they also appear in our less structured treatment (Unrestricted Migration). Most likely, the diversity results partly from the presence of small fitness differences between genotypes, such that new beneficial mutations may arise before old beneficial mutations fix (leading to a form of clonal interference). We also note that the Unrestricted treatment possessed some degree of population structure (i.e., this was not a “well-mixed” population), which could also contribute to diversity. Such diversity may enable metapopulations in the Unrestricted treatment to explore multiple domains simultaneously. However, the main effects of structure outlined in this paper still apply. If the landscape is multipeaked, metapopulations in the Restricted treatment are expected to sample *a greater number of domains* (and indeed, the Restricted metapopulations had significantly greater diversity). We emphasize that the same violations of SSWM assumptions occur in the NK model (see Supplemental Figure 1 for diversity profiles), in which the same migration treatments were used, and we note that the Tortoise-Hare pattern is observed for sufficiently rugged landscapes (Figure 2). Thus, even in populations violating strict SSWM assumptions, the greater parallel search that comes with greater population structure is predicted to lead to better long-term adaptation in rugged landscapes with heterogeneity in peak height.

Full Data, Simulation Code, and Statistical Scripts are available on the Kerr Lab Wiki: http://kerrlab.org/Public/RugLand

